# An optimized acetylcholine sensor for monitoring *in vivo* cholinergic activity

**DOI:** 10.1101/861690

**Authors:** Miao Jing, Yuexuan Li, Jianzhi Zeng, Pengcheng Huang, Miguel Skirzewski, Ornela Kljakic, Wanling Peng, Tongrui Qian, Ke Tan, Runlong Wu, Shichen Zhang, Sunlei Pan, Min Xu, Haohong Li, Lisa M. Saksida, Vania F. Prado, Tim Bussey, Marco A.M. Prado, Liangyi Chen, Heping Cheng, Yulong Li

## Abstract

The ability to directly measure acetylcholine (ACh) release is an essential first step towards understanding its physiological function. Here we optimized the GRAB_ACh_ (<u>G</u>PC<u>R</u>-<u>A</u>ctivation–<u>B</u>ased-<u>ACh</u>) sensor with significantly improved sensitivity and minimal downstream coupling. Using this sensor, we measured *in-vivo* cholinergic activity in both *Drosophila* and mice, revealing compartmental ACh signals in fly olfactory center and single-trial ACh dynamics in multiple regions of the mice brain under a variety of different behaviors

---

Cholinergic signals mediated by the neurotransmitter ACh are involved in a wide range of physiological processes, including muscle contraction, cardiovascular function, neural plasticity, attention and memory^1-3^. Previously, cholinergic activity was mainly measured using either electrophysiology to record nicotinic receptor-mediated currents^4, 5^ or microdialysis followed by biochemical purification and identification^6^. However, these methods generally lack both cell-type specificity and the spatial-temporal resolution needed to precisely dissect cholinergic signals *in vivo*. Combining the type 3 muscarinic ACh receptor (M_3_R) with the conformational-sensitive circular permutated GFP (cpGFP), we recently developed GACh2.0 (short as ACh2.0), a genetically encoded GRAB (<u>G</u>PC<u>R</u>-<u>A</u>ctivation <u>B</u>ased) ACh sensor that can convert the ACh-induced conformational change on M_3_R into a sensitive fluorescence response^7^. The ACh2.0 sensor responds selectively to physiological concentration of ACh with an EC_50_ of 2 μM and has been used in several model organisms to detect the endogenous release and regulation of cholinergic signals. Here, we optimized the GRAB_ACh_ sensor using site-directed mutagenesis and cell-based screening to further increase the sensitivity.

To improve the performance of the GRAB_ACh_ sensor, we focused on the interface between M_3_R and cpGFP, including the receptor’s third intracellular loop (ICL3) and linker peptides, as well as critical residues in cpGFP that contribute to its fluorescence intensity (Figs. 1A and S1A-D). Our initial screening based on medium-throughput imaging identified several variants with improved performance; these variants were subsequently verified using confocal microscopy (see Methods for details). The sensor with the largest ACh-induced fluorescence response was selected for further study and is named as GRAB_ACh3.0_ or ACh3.0 (Fig. 1A). We also generated a ligand-insensitive form of ACh3.0 by introducing the W200A mutation into the receptor^8^ (Figs. 1A and S1E). When expressed in HEK293T cells or cultured neurons, the ACh3.0 sensor localized to the plasma membrane of the soma, and trafficked to dendrites and axons in neurons (Fig. 1B-D). Moreover, compared to ACh2.0, the ACh3.0 sensor had a significantly larger fluorescence change (ΔF/F_0_∼280%) in response to 100 μM ACh (Figs. 1B-D and S2A-E); in contrast, the ligand-insensitive ACh3.0-mut sensor had no detectable fluorescence change, even at high ACh concentrations (e.g. 100 μM) (Figs. 1B-D and S1F). Importantly, ACh3.0 shared a similar affinity for ACh as ACh2.0 (EC_50_∼2 μM) and did not respond to other major neurotransmitters (Fig. 1E, F). Moreover, the ACh-induced response in ACh3.0 could be blocked by the muscarinic receptor antagonist tiotropium (Tio) (Fig. 1F). Using multiple assays including intracellular Ca^2+^ imaging, a Gq-dependent luciferase complementary assay^9^, and a β-arrestin–dependent TANGO assay^10^, we confirmed that ACh3.0 has virtually no coupling with major GPCR-mediated downstream pathways (Figs. 1G,H and S3A-E).

**Figure 1:**
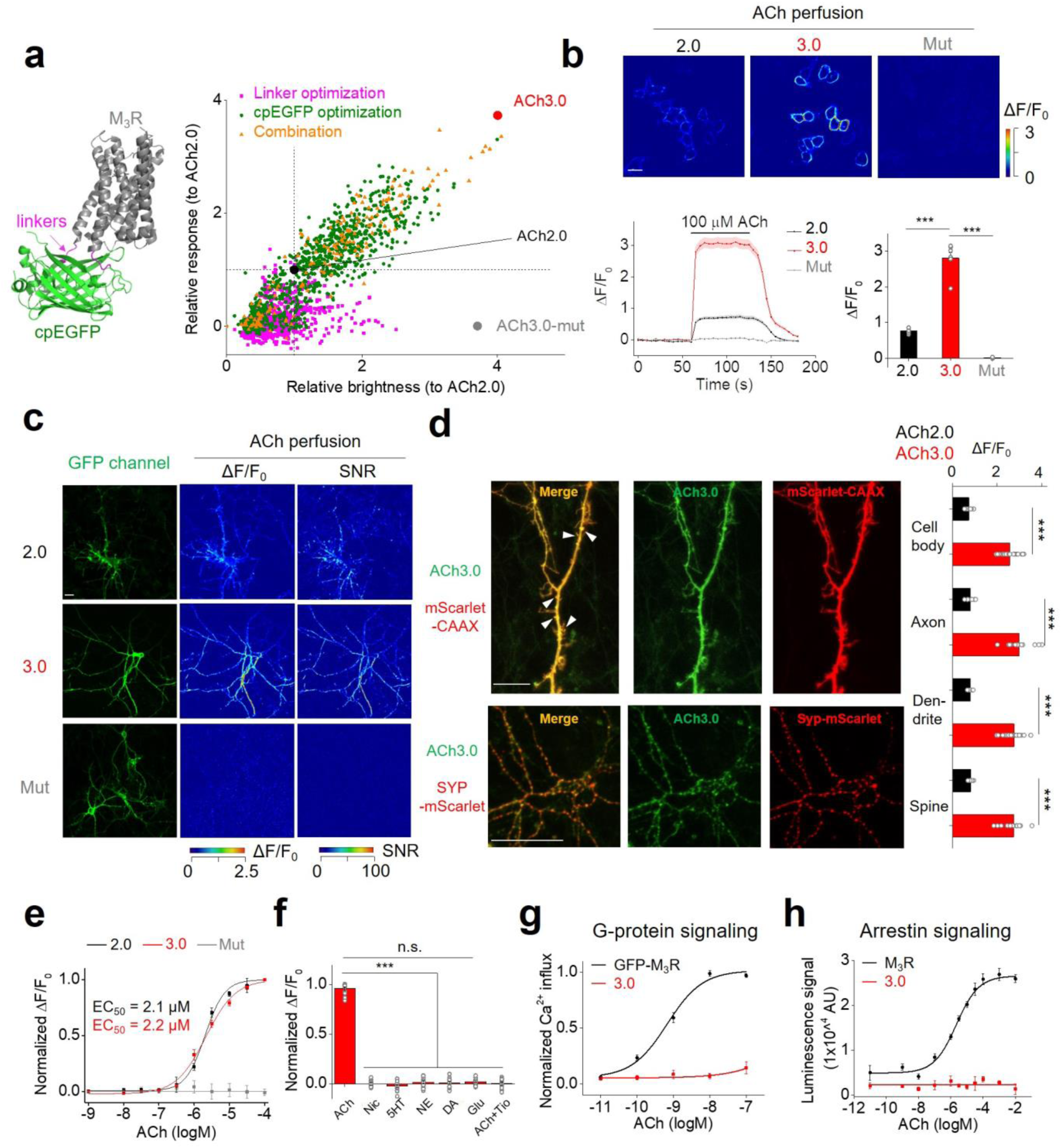
Optimization and *in vitro* characterization of next-generation GRAB_ACh_ sensors. **A:** Left: cartoon illustration showing the predicted structure of GRAB_ACh_ sensors. Right: site-directed random mutagenesis is performed in the linkers between the receptor and cpEGFP (magenta), cpEGFP (green), or both (orange), and the performance of each variant expressed in HEK293T cells is calculated relative to ACh2.0 (black) and plotted. The final optimized sensor, ACh3.0 is indicated in red, and the ligand-insensitive ACh3.0-mut sensor (with W200A mutation) is indicated in gray. All the response and brightness of candidates are measured using Opera Phenix high content imaging system (See Methods for detail). Each data point represents the average response measured in >100 cells per candidate. The response and brightness of ACh2.0 are calculated from confocal imaging and used for normalization. The crystal structures are adopted from PDB database (4DAJ for M_3_R and 3EK4 for cpGFP). **B:** The performance of the ACh2.0, ACh3.0, and ACh3.0-mut sensors expressed in HEK293T cells in response to 100 μM ACh in confocal imaging. Top: pseudocolor images of the peak response (ΔF/F_0_) in the presence of 100 μM ACh. Bottom: representative traces and group data; n=5, 7, and 4 coverslips for ACh2.0, ACh3.0, and ACh3.0-mut, respectively, with an average of >20 cells per coverslip. **C:** The performance of the ACh2.0, ACh3.0 and ACh3.0-mut sensors expressed in cultured neurons in response to 100 μM ACh. The raw GFP fluorescence and pseudocolor images of the peak response to ACh (ΔF/F_0_) and signal-to-noise ratio (SNR) are shown. **D:** Left: representative images of ACh3.0 expressed together with subcellular markers fused to the red fluorescent protein mScarlet. Dendritic spines are indicated by white arrowheads. Right: group data summarizing the fluorescence response of ACh2.0 (black bars) and ACh3.0 (red bars) to 100 μM ACh measured in the indicated neuronal compartments (cell body: n=11 for ACh2.0 and n=18 for ACh3.0; axon: n=8 for ACh2.0 and n=14 for ACh3.0; dendrite: n=18 for ACh2.0 and n=26 for ACh3.0; spine: n=10 for ACh2.0 and n=16 for ACh3.0). **E:** Dose-response curves for the fluorescence response of ACh2.0 (black), ACh3.0 (red), and ACh3.0-mut (gray) to ACh, with corresponding EC_50_ values (n=11,12 and 16 neurons for ACh2.0, ACh3.0 and ACh3.0-mut, respectively). **F:** The fluorescence response of ACh3.0 to the indicated compounds (n=12 neurons each). ACh: 100 μM; nicotine (Nic): 50 μM; 5-HT: 1 μM; norepinephrine (NE): 10 μM; dopamine (DA): 20 μM; glutamate (Glu): 10 μM; and tiotropium (Tio): 2 μM. **G:** The normalized Ca^2+^ response to ACh in HEK293T cells expressing GFP-M_3_R or ACh3.0 (n=22 and 15 cells for GFP-M_3_R and ACh3.0, respectively). **H:** The β-arrestin dependent luminescence signal in HEK293T cells expressing GFP-M_3_R or ACh3.0 in response to ACh at indicated concentration (n>100 cells/well from n=4 wells). Scale bars represent 10 μm. \*\*\**p*<0.001 and n.s., not significant.

We previously reported that the ACh2.0 sensor could be used to record endogenous ACh release, including the GABA_B_R-dependent potentiation of ACh release in the MHb-IPN (medial habenula-interpeduncular nucleus) projection in acute mouse brain slices^7, 11^. We next tested whether ACh3.0 had better performance in reporting endogenous ACh in MHb-IPN slices. Consistent with our *in vitro* results, the ACh3.0 sensor has a significantly larger fluorescence increase compared to ACh2.0 in response to high-frequency (>10 Hz) electrical stimulation of the cholinergic fibers, both in control ACSF solution and in the presence of the GABA_B_R agonist baclofen (Bac) (Figs. 2B, C and S4A-F). Application of the GABA_B_R antagonist saclofen (Sac) reversed the baclofen-induced potentiation, and further adding Tio eliminated the stimulation-evoked response (Fig. 2D). Moreover, the electrical stimulation-evoked signal was increased by the potassium channel blocker 4-AP and eliminated by Cd^2+^, consistent with Ca^2+^-dependent ACh release (Figs. 2E and S4G, H). Using a brief (100-ms) electrical stimulation, we then measured the kinetics of the fluorescence response, yielding tau_on_ and tau_off_ values of ∼105 ms and 3.7 s, respectively (Fig. 2F).

**Figure 2:**
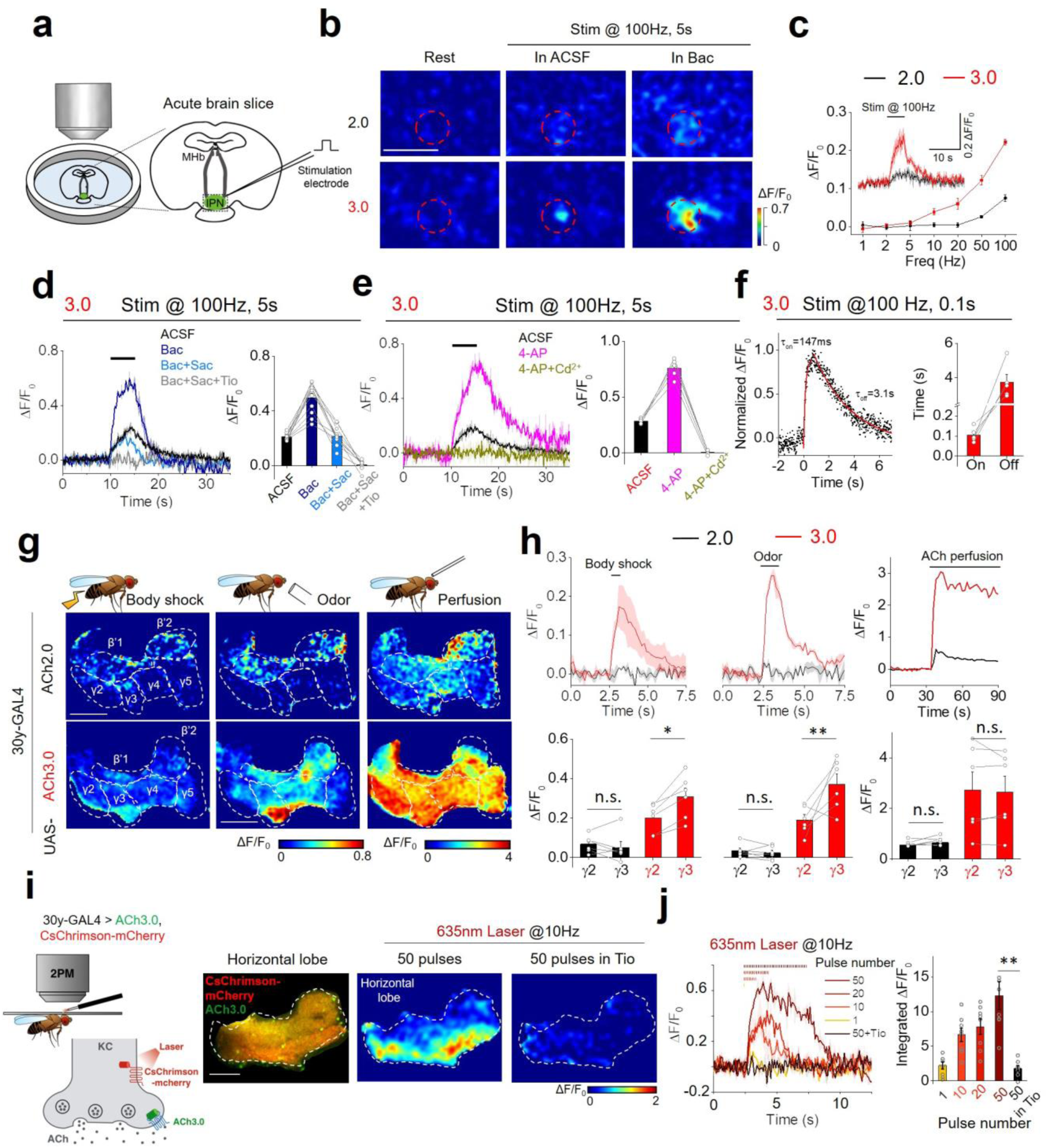
Probing ACh dynamics in acute mouse brain slices and *in vivo* in *Drosophila*. **A:** Schematic illustration depicting the two-photon imaging of acute MHb-IPN brain slices prepared from mice expressing ACh2.0 or ACh3.0 in the IPN. A bipolar electrode placed in the IPN is used to evoke endogenous ACh release. **B:** Pseudocolor images of the fluorescence response (ΔF/F_0_) of ACh2.0 and ACh3.0 to electrical stimuli (100 Hz for 5 s) in ACSF or 2 μM baclofen (Bac). The red dashed circles indicate the regions of interest (30 μm in diameter) used for quantification. Data are representative of 5-10 slices from 3-7 mice. **C:** Group summary of the fluorescence response of ACh2.0 and ACh3.0 to electrical stimuli at the indicated frequencies (n=11 slices from 8 mice). The inset shows representative traces of ACh2.0 and ACh3.0 in response to 100-Hz electrical stimulation. **D:** Representative traces and group summary of the fluorescence response of ACh3.0 to electrical stimulation in either ACSF or the indicated drugs; n=5 slices from 5 mice per group. Baclofen (Bac): 2 μM; saclofen (Sac): 100 μM; and tiotropium (Tio): 10 μM. **E:** Representative traces and group summary of the fluorescence response of ACh3.0 to electrical stimulation in ACSF, 4-AP (100 μM), or 4-AP with Cd^2+^(100 μM); n=5 slices from 5 mice per group. **F:** Fluorescence traces of ACh3.0 in response to 100-ms electrical stimulation. The rising and decay phases of the fluorescence signals are fitted to a single-exponential function, and the time constants are summarized on the right; n=5 slices from 5 mice per group. **G:** Pseudocolor images of the fluorescence response in the mushroom body horizontal lobe in transgenic flies expressing ACh2.0 or ACh3.0 during body shock (left), odorant application (middle), and exogenous ACh perfusion (right). **H:** Top: fluorescence traces measured in the mushroom body in transgenic flies expressing ACh2.0 (black) or ACh3.0 (red); where indicated, body shock, odorant stimulation, or ACh is applied. Bottom: group summary of the fluorescence responses measured in the γ2 and γ3 lobes in the mushroom body; n=6 flies per group. **I:** Left: schematic illustration depicting the experimental setup; CsChrimson-mCherry and ACh3.0 sensors are expressed in Kenyon cells (KCs) in the mushroom body, and 635-nm laser light is used to activate cholinergic KCs. Right: fluorescence images and pseudocolor images of ACh3.0 sensors in response to 635-nm laser stimulation in the absence or presence of Tio (10 μM). **J:** Representative traces and group summary of the fluorescence response of ACh3.0 to the indicated number of 635-nm laser pulses applied at 10 Hz; n=8 flies per group. Scale bars represent 50 μm (B) and 25 μm (G and I). \**p*<0.05, \*\**p*<0.01, and n.s., not significant.

Next, we used *in vivo* two-photon imaging to compare the performance of ACh3.0 and ACh2.0 in transgenic *Drosophila* expressing the ACh sensors in Kenyon cells (KCs) of the olfactory mushroom body (Fig. 2G). Physiologically relevant stimuli, including body shock to the abdomen and odor stimulation, elicited only a minor fluorescence signal in the horizontal lobe of the mushroom body in ACh2.0-expressing flies. In contrast, the same stimuli induced a significantly larger response in ACh3.0-expressing flies, and such increase was higher in γ3 lob compared with adjacent γ2 lobe, which revealed compartment-specific release of ACh (Fig. 2G, H). In addition, we monitored ACh dynamics in response to direct neuronal activation of KCs via CsChrimson-mediated optogenetics^12^ or electrical stimulation. A single 635-nm laser pulse evoked a clear increase in ACh3.0 fluorescence, and multiple pulses applied at 10-Hz induced a progressively larger response that was largely eliminated by Tio application (Fig. 2I, J). Electrical stimulation of KCs confirmed the frequency-dependent fluorescence increase in the ACh3.0 sensor, with rapid kinetics (tau_on_ ∼0.09 s and tau_0ff_ ∼0.91 s) (Fig. S5). Taken together, these data show that the ACh3.0 sensor can reliably detect ACh release with high sensitivity and spatial-temporal resolution.

Cholinergic signaling plays a key role in modulating a variety of physiological processes, including plasticity and arousal. We therefore examined whether ACh3.0 can be used *in vivo* to monitor ACh release in behaving mice. In the mouse brain, basal forebrain cholinergic neurons project to the amygdala, hippocampus and cortical regions^13^. We measured foot-shock evoked cholinergic signal in the basolateral amygdala (BLA), which is believed to be important in aversive associate learning^14^. A brief foot-shock stimulus induced a reproducible and time-locked increase in ACh3.0 fluorescence (Fig. 3A, B). We then combined pharmacology and genetics to confirm the signal’s specificity. Treating the mice with the acetylcholinesterase inhibitor (AChEI) donepezil (Done) significantly slowed the fluorescence decay, while the M_3_R antagonist scopolamine (Scop) abolished the foot-shock-induced fluorescence response. As a control, no response was observed in mice expressing ACh3.0-mut (Fig. 3D) or in mice lacking the vesicular ACh transporter (VAChT)^15^, suggesting the synaptic origin of ACh release (Figs. 3E and S6A-C). Taken together, these *in vivo* data indicates that the ACh3.0 sensor can reliably report foot-shock induced cholinergic signaling in the BLA. Next, we examined whether ACh3.0 could stably report ACh signaling over a longer time period by recording ACh dynamics in the mouse hippocampus during the sleep-wake cycle. A simultaneous recording of EEG and EMG was used to monitor the animal’s sleep status (Fig. 3F, see Methods for detail). We found that the average fluorescence signal was larger in wakefulness and REM sleep, but lower during non-REM sleep (Figs. 3G, H), consistent with our previous optrode-based recordings of the firing rate of basal forebrain cholinergic neurons^16^. Interestingly, ACh3.0 was also able to detect ACh increase induced by micro-arousal events during NREM sleep (Fig. 3G). Overall, these data illustrate that the ACh3.0 has both the high sensitivity, rapid kinetics and stability needed to reliably track long-term physiological cholinergic signal *in vivo*.

**Figure 3:**
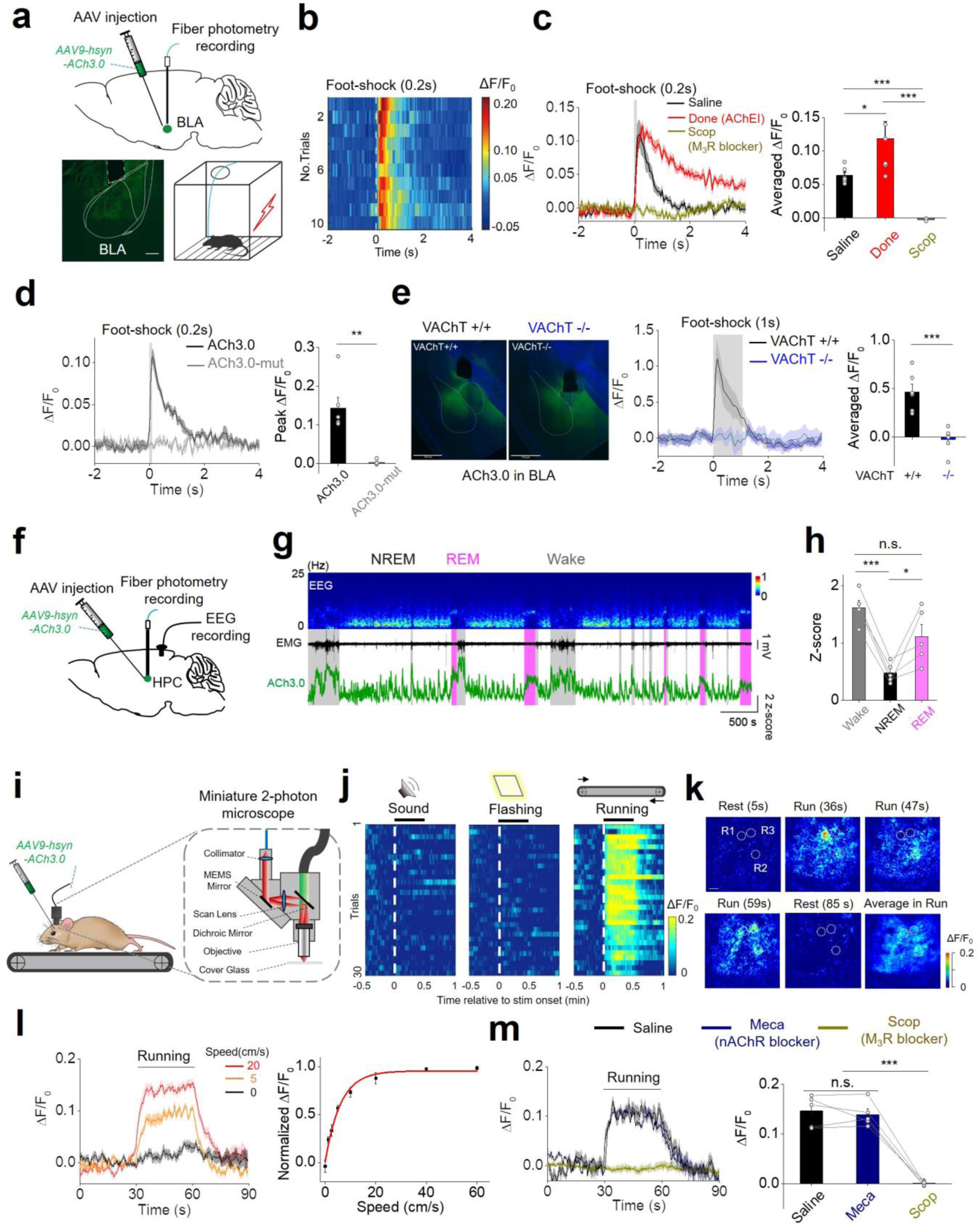
Monitoring *in vivo* ACh dynamics in mice. **A:** Top: schematic diagram depicting the injection of an AAV encoding the ACh3.0 sensor into the basolateral amygdala (BLA); the fluorescence response is recorded using fiber-photometry. Bottom: fluorescence of the ACh3.0 sensor expressed in the BLA (left) and a cartoon illustration of the foot-shock experiments (right). **B:** Pseudocolor fluorescence responses of ACh3.0 in the BLA to ten 0.2-s foot-shock stimuli at 0.4 mA. **C:** Representative traces and group summary of the fluorescence response of ACh3.0 in the BLA of mice following an i.p. injection of saline (black), the acetylcholinesterase inhibitor donepezil (Done, red, 3 mg/kg body weight), or the M_3_R antagonist scopolamine (Scop, gray, 6 mg/kg body weight). The average fluorescence response is calculated using the 1-s mean fluorescence after the initiation of foot-shock (n=6 mice per group). **D:** Similar to C, showing the fluorescence response of ACh3.0 and ACh3.0-mut to a 0.2-s foot-shock; n=6 and 4 mice for ACh3.0 and ACh3.0-mut, respectively. **E:** Left: fluorescence images of ACh3.0 expressed in the BLA of control mice (VAChT^+/+^, black) and VAChT forebrain knockout mice (VAChT^-/-^, blue). Middle and right: representative traces and group summary of the response measured in the BLA of VAChT^+/+^ and VAChT^-/-^ mice to 1-s foot-shock; n=6 mice per group. Scale bars represent 750 μm. **F:** Schematic diagram depicting the injection of an AAV expressing ACh3.0 into the mouse hippocampus (HPC); fluorescence is recorded in the mice during the sleep-wake cycle. The placement of intracranial EEG recording electrodes is also indicated. **G:** Representative recording of EEG (top), EMG (middle), and ACh3.0 fluorescence (bottom) in a mouse during the sleep-wake cycle. The mouse’s sleep/wake status (wake, NREM sleep, or REM sleep) is determined using the EEG and EMG data and is indicated. **H:** Group summary of the ACh3.0 fluorescence response (expressed as a Z-score) in mice while awake and during NREM and REM sleep; n=5 mice per group. **I:** Schematic illustration depicting the experiment in which mice expressing ACh3.0 in the visual cortex are placed on a treadmill and ACh3.0 fluorescence is recorded using the miniature two-photon microscope (shown in detail on the right). **J:** Pseudocolor fluorescence responses of ACh3.0 to an auditory stimulus (30 s of a 7000-Hz tone, left), a visual stimulus (30-s of flashing light at 2 Hz, middle), or running on the treadmill (right). The responses of 30 consecutive trials are recorded and are plotted relative to the onset of each stimulus. **K:** Pseudocolor images showing the spatial-temporal distribution of ACh3.0 fluorescence during locomotion from a single trial. R1, R2 and R3 are three representative ROIs (40 μm in diameter) indicating spatial selective ACh signals at indicated time point during running. The averaged fluorescent signal during entire running process is also shown. Scale bar, 50 μm. **L:** Representative traces and group summary of ACh3.0 fluorescence in mice recorded while running on a treadmill at the indicated speeds; each trace is averaged from 10 trials; n=5 mice. **M:** Representative traces and group summary of ACh3.0 fluorescence measured in mice while performing the running task; where indicated, the mice receive an i.p. injection of saline (black), the nAChR blocker mecamylamine (Meca, 2 mg/kg body weight, blue), or the M_3_R antagonist scopolamine (Scop, 20 mg/kg body weight, dark yellow); each trace is averaged from 10 trials; n=5 mice per group. \**p*<0.05, \*\**p*<0.01, \*\*\**p*<0.001, and n.s., not significant.

Next, to demonstrate that the applicability of using the ACh3.0 sensor in tracking cholinergic signaling with high spatial resolution in freely behaving animals, we measured cortical ACh signals in response to different stimuli using miniature two-photon microscopy^17^. We expressed the ACh3.0 sensor in the mouse visual cortex and monitored the fluorescence signal using a miniature two-photon microscope attached to the head of the mouse (Fig. 3I). During two-photon imaging, we placed the mouse on a treadmill and measured the ACh3.0 signal in response to auditory or visual stimuli or during animal’s locomotion. We observed a robust increase in ACh3.0 fluorescence when the mouse was running, but not during the application of sensory stimulation (Fig. 3J). We could also reveal spatial selective ACh signals at certain time points during running from a single imaging trial, indicating the ACh3.0 has high spatial and temporal resolution to resolve ACh patterns (Fig. 3K). Moreover, the fluorescence response positively correlated with the running speed of the mice (Fig. 3L). The running-related increase in the ACh3.0 signal was abolished by M_3_R antagonist Scop, but not nAChR blocker mecamylamine (Meca), indicating a high specificity of the measured fluorescence signal (Fig. 3M).

Our previous ACh2.0 sensor has been reported to couple with downstream Gq signaling, although with ∼10 fold less affinity compared with native M_3_R. In contrast, the optimized ACh3.0 sensor has negligible coupling to G protein signaling in two independent assays (Figs. 1G and S3D). We reasoned that these improvements might be due to truncation of the receptor’s ICL3 domain, which prevents the receptor from interacting with the G protein and β-arrestin. Thus, the ACh3.0 sensor can be viewed as a highly sensitive ACh detector that functions independently from cellular signaling. Indeed, we confirmed that expressing ACh3.0 does not affect the physiological properties of neurons, as odorant-evoked Ca^2+^ transients in *Drosophila* expressing both ACh3.0 and jRCaMP1a were similar to those in flies expressing only jRCaMP1a (Fig. S3F, G).

In conclusion, we engineered the GRAB_ACh3.0_ with higher sensitivity while maintained its specificity, precise temporal and spatial resolution and high photo-stability in detecting ACh. The new sensor provides a robust tool for testing previously suggested hypotheses on the dynamics of cholinergic activity under both physiological and pathophysiological conditions. With respect to studying physiological processes, the ACh3.0 sensor allows the real-time visualization of compartment-specific ACh release in the *Drosophila* olfactory system and cholinergic dynamics during the sleep-wake cycle in mice, shedding new light on how cholinergic signaling is regulated. The combination of improved ACh sensor with advanced imaging techniques will help to address fundamental biological questions in the future.

## Methods

### Animals

Male and female P0 Sprague–Dawley rats were used to prepare cultured cortical neurons; P28-48 wild-type C57BL/6N mice were used to prepare the acute brain slices and two-photon *in vivo* imaging. C57BL/6J mice were used for fiber photometry recording. Mice lacking the vesicular acetylcholine transporter in the forebrain were generated as previously described^18^ by crossing VAChT^flox/flox^ mice (Chat/Slc18a3^tm1.2Vpra^ generated in a mixed C57BL/6J × 129/SvEv background, backcrossed to C57BL/6J for 10 generations)^15^ with Nkx2.1-Cre mice (The Jackson Laboratory, stock no. JAX008661), yielding VAChT^-/-^ offspring and control (VAChT^flox/flox^) littermates (referred to here as VAChT^+/+^). All rodents were either family-housed or pair-housed in a temperature-controlled room with a 12-h/12-h light/dark cycle. All procedures for animal surgery and experimentation were performed using protocols approved by the Animal Care & Use Committees at the Chinese Institute for Brain Research, the Peking University, the Chinese Academy of Sciences (CAS), Huazhong University of Science and Technology, and the University of Western Ontario (2016-104), and were performed in accordance with the guidelines established by US National Institutes of Health. To generate transgenic *Drosophila melanogaster*, the two plasmids 10xUAS-IVS-ACh3.0-p10 and 10xLexAop2-IVS-ACh3.0-p10 were constructed and integrated into attp40 or VK00005 of fly genome mediated by PhiC31. The embryo injections were performed at Core Facility of *Drosophila* Resource and Technology, Shanghai Institute of Biochemistry and Cell Biology, CAS. Transgenic flies were raised on conventional corn meal at 25°C, with ∼70% humidity, under 12-h/12-h light/dark cycle. The fly lines used in this study: 30y-GAL4 (BDSC: 30818) from Yi Rao (Peking University), UAS-CsChrimson-mcherry and UAS-jRCaMP1a (BDSC: 64427) from Chuan Zhou (Institute of Zoology, CAS), and MB247-LexA from Yi Zhong (Tsinghua University).

### Molecular biology

Plasmids were generated using the Gibson assembly method^19^. DNA fragments were generated using PCR amplification using primers (Thermo Fisher Scientific) with 30-bp overlap. The fragments were then assembled using T5-exonuclease (New England Biolabs), Phusion DNA polymerase (Thermo Fisher Scientific), and Taq ligase (iCloning). All sequences were verified using Sanger sequencing at the Sequencing Platform in the School of Life Sciences of Peking University. For screening in HEK293T cells, the ACh sensor constructs were cloned in the pDisplay vector (Invitrogen) followed by the IRES-mCherry-CAAX sequence, which served as a membrane marker to calibrate the signal intensity. Site-directed mutagenesis of the linker sequences and residues in cpEGFP was performed using primers containing randomized NNB codons (48 codons in total, encoding all 20 amino acids; Thermo Fisher Scientific). For AAV package, the ACh3.0 sensor and mutant ACh3.0-mut sensor were cloned into an AAV vector under the control of the human synapsin promoter. To generate transgenic *Drosophila* lines, the ACh3.0 sensor was cloned into the pJFRC28 vector, which was then used to generate transgenic flies via PhiC31-mediated site-directed integration into attp40.

### Cell culture, transfection, and imaging

HEK293T cells (ATCC cell line CRL-3216) were cultured at 37°C in 5% CO_2_ in DMEM (GIBCO) supplemented with 10% (v/v) fetal bovine serum (GIBCO) and 1% penicillin-streptomycin (GIBCO). HEK293 cells stably expressing a tTA-dependent luciferase reporter and a β-arrestin2-TEV fusion construct were a gift from Bryan L. Roth. Rat cortical neurons were prepared from P0 Sprague–Dawley rat pups (both male and female; Beijing Vital River). In brief, the brains were dissected, and cortical neurons were dissociated in 0.25% Trypsin-EDTA (GIBCO), plated on 12-mm glass coverslips coated with poly-D-lysine (Sigma-Aldrich), and cultured at 37°C in 5% CO_2_ in neurobasal medium containing 2% B-27 supplement, 1% GlutaMax, and 1% penicillin-streptomycin (all from GIBCO). HEK293T cells were transfected using the polyethylenimine (PEI) method (with a typical ratio of 1 µg DNA to 4 µg PEI); the media was replaced 4–6 h later, and cells were imaged 24 h after transfection. Cultured neurons were transfected at 7–9 days *in vitro* using the calcium phosphate transfection method, and experiments were performed 48 h after transfection. For screening candidate sensors, cultured HEK293T cells expressing the various candidate sensors were first imaged using the Opera Phenix High-Content Screening System (PerkinElmer) equipped with a 60 × /1.15 NA water-immersion objective, a 488-nm laser, and a 561-nm laser; the ACh sensor’s signal was obtained using a 525/50-nm emission filter, and the mCherry-CAAX signal was obtained using a 600/30-nm emission filter. The ratio between green (G) and red (R) fluorescence was calculated before and after application of 100 μM ACh, and the change in the G/R ratio was used as the fluorescence response; the peak G/R ratio was used as a brightness index. Candidate sensors with the best performance were subsequently imaged using a Ti-E A1 inverted confocal microscope (Nikon) equipped with a 40 × /1.35 NA oil immersion objective, a 488-nm laser, and a 561-nm laser. Drugs were prepared in Tyrode’s solution containing (in mM): 150 NaCl, 4 KCl, 2 MgCl_2_, 2 CaCl_2_, 10 HEPES, and 10 glucose (pH 7.4) and perfused into the imaging chamber. The ACh sensor’s signal was obtained using a 525/50-nm emission filter, and the mCherry signal was obtained using a 595/50-nm emission filter. Cultured neurons expressing the ACh sensor were similarly imaged using the inverted confocal microscope with drugs added by perfusion.

### Slice preparation and imaging

Adeno-associated viruses (AAVs) expressing either ACh2.0 or ACh3.0 were packaged by Vigene Biosciences, and were injected into the mouse interpeduncular nucleus (IPN) (500 nl per mice, titer: 1×10^13 vg/ml). Two weeks after virus injection, the animals were anesthetized with an i.p. injection of Avertin (250 mg/kg body weight), and the heart was perfused with 5ml slicing buffer containing (in mM): 110 choline-Cl, 2.5 KCl, 1.25 NaH_2_PO_4_, 25 NaHCO_3_, 7 MgCl_2_, 25 glucose, and 2 CaCl_2_. The mice were then decapitated and brains were removed immediately and placed directly in cold oxygenated slicing buffer. The brains were first blocked at an ∼45° angle relative to the horizontal plane and then sectioned into 250-μm thick slices using a VT1200 vibratome (Leica); the sections were transferred to oxygenated Ringer’s buffer containing (in mM): 125 NaCl, 2.5 KCl, 1.25 NaH_2_PO_4_, 25 NaHCO_3_, 1.3 MgCl_2_, 25 glucose, and 2 CaCl_2_. The slices were then recovered at 34°C for at least 40 min. For two-photon fluorescence imaging, slices were transferred to an imaging chamber and placed in an FV1000MPE two-photon microscope (Olympus) equipped with a 40 × /0.80 NA water-immersion objective and a mode-locked Mai Tai Ti:Sapphire laser (Spectra-Physics) tuned to 920 nm for excitation and a 495∼540-nm filter for signal collection. For electrical stimulation, a bipolar electrode (cat#WE30031.0A3, MicroProbes) was positioned near the IPN region under the fluorescence guidance, and the imaging and stimulation were synchronized using an Arduino board with a custom program. The stimulation voltage was set at ∼4 V, and the duration of each stimulation pulse was set at 1 ms. Drugs were added by perfusion or were bath-applied.

### Two-photon imaging in *Drosophila*

Female *Drosophil*a *melanogaster* within 3 weeks after eclosion were used for imaging experiments. The fly was mounted on a customized chamber by tape, in a way the antenna and abdomen exposed to the air. The cuticle between compound eyes, as well as air sacs and fat bodies were removed to expose the brain which was then bathed in the saline, i.e. adult hemolymph-like solution (AHLS): (in mM) 108 NaCl, 5 KCl, 5 HEPES, 5 Trehalose, 5 sucrose, 26 NaHCO_3_, 1 NaH_2_PO_4_, 2 CaCl_2_ and 2 MgCl_2_. The same Olympus two-photon microscope as well as electrical stimulation equipment used for brain slices imaging was also used here. 920 nm laser was used for excitation. 495∼540 nm filter was used for ACh2.0 or ACh3.0 imaging, and 575∼630 nm filter was used for jRCaMP1a imaging. For odor stimulation, the odorant isoamyl acetate (Sigma-Aldrich, Cat#306967) was first diluted by 200-fold in mineral oil in a bottle and second diluted by 5-fold in air, which was then delivered to the fly’s antenna at a rate of 1000 ml/min. For body shock, two wires were attached to the abdomen of the fly, and a 60∼80 V electrical pulse were delivered for 500 ms during stimulation. For ACh application, a patch of blood-brain-barrier of the fly was carefully removed by tweezers before imaging, and the saline containing ACh was delivered to the brain to 20 mM final concentration. For optogenetic stimulation, the 635 nm laser (1 ms pulses at 10 Hz) were delivered through optical fibers placed near the fly brain. Flies were fed with corn meal containing 10 μM β-carotene (Sigma-Aldrich) 3 days before optogenetics experiments. For electrical stimulation, a glass electrode (resistance ∼0.2 MΩ) was placed in the region of the Mushroom Bodies and the stimulation voltage was set at 20∼80 V. Tiotropium was added directly to the saline to the final concentration, and the following experiments were performed 10 min after application. The nicotinic receptor blocker mecamylamine (100 μM) was bath applied in electrical stimulation and optogenetic stimulation experiments. The sampling rates of imaging were set at 7 Hz during odor or body shock stimulation, 0.5 Hz during ACh application, 10 Hz during optogenetic stimulation and 12 Hz during electrical stimulation. Arduino was used to synchronized stimulation delivery and imaging with a custom program.

### Fiber-photometry, EEG and EMG recordings in mice

The AAV carrying ACh3.0 were injected into the basolateral amygdala (AP −1.4 mm, ML 3.1 mm, DV 4.1 mm) by a microsyringe pump, or into the hippocampus dorsal CA1 (AP −2.2 mm, ML 1.5 mm, DV 1.2 mm) using Nanoject II (Drummond Scientific) via a glass pipette. An optic fiber (230 µm, 0.37NA or 400 μm, 0.48 NA for BLA recording; 200 µm, 0.39NA for hippocampus recording) was inserted into the same coordinate used for virus injection. For the recording in BLA, the f-scope fiber photometry system (BiolinkOptics, China) with laser power adjust at 20-30 μW, or the fiber photometry system from Doric Lense (Quebec, Canada) with 465nm LED power set at 20-25 μW were used to record fluorescence signals. For recording in the hippocampus, an optic fiber (Thorlabs, FT200UMT) was attached to the implanted ferrule via a ceramic sleeve and recorded emission fluorescence using fiber photometry. The photometry rig was constructed using parts from Doric Lens,including a fluorescence optical mini cube (FMC4_AE(405)_E(460-490)_F(500-550)_S), a blue LED (CLED_465), a LED driver (LED_2) and a photo receiver (NPM_2151_FOA_FC). Data collected by the Quebec system at 186 Hz were first filtered at 6-Hz low-pass and analyzed. Data from the hippocampus recording were first binned into 1Hz, subtracted the background autofluorescence and then analyzed.

To implant EEG and EMG recording electrodes, mice were anesthetized with isoflurane (5% induction; 1.5 - 2% maintenance) and placed on a stereotaxic frame with a heating pad. Two stainless steel screws for EEG were inserted into the skull above the visual cortex, two other screws were inserted into the skull above the frontal cortex, two insulated EMG electrodes were inserted into the neck muscle, and reference electrode was attached to a screw inserted into the skull above the cerebellum. The implant was secured to the skull with dental cement. All the experiments were carried out at least one week after the surgery. The EEG/EMG signals were recorded using TDT system-3 amplifiers (RZ2 + PZ5) with a high-pass filter at 0.5 Hz and digitized at 1500 Hz. Spectral analysis was carried out using fast Fourier transform (FFT) with a frequency resolution of 0.18 Hz. The brain states were scored every 5 seconds semi-automatically using a MATLAB GUI and validated manually by trained experimenters. Wakefulness was defined as desynchronized EEG and high EMG activity; NREM sleep was defined as synchronized EEG with high-amplitude delta activity (0.5 - 4 Hz) and low EMG activity; REM sleep was defined as high power at theta frequencies (6 - 9 Hz) and low EMG activity.

For ACh recording during foot-shock, mice were first habituated to the behavioral chamber set at 65db background noise and under infrared light for video visualization for 4 sessions (10 min per session, twice/day). Then, mice were habituated for 5 min before receiving foot-shock stimuli (10 trials of 0.2s or 1s long, 0.4 mA foot-shock intensity, 20s inter-trial intervals). Each foot-shock was coupled to the delivery of a 10ms long TTL output for synchronization with the fiber photometry recordings.

### Two-photon imaging in mice

To express ACh3.0, the mice were initially anesthetized with an injection of Avertin, the skin was retracted from the head, and a metal recording chamber was affixed. On the second day, the mice were anesthetized again, the skull above the visual cortex was opened, and ∼500 nl of AAV was injected at the depth of 0.5 mm. A 4 mm x 4 mm square coverslip was used to replace the skull. Two-photon imaging in the visual cortex was performed 3 weeks after virus injection as previously reported^17^. In brief, the mice were first attached with the baseplate and allowed to habituate for 2-3 days before the experiment. During the experiment, the miniature two-photon microscope was placed on the baseplate, and the mice were head-fixed on a treadmill controlled by an electric motor. The output trigger from the computer was used to synchronize the imaging with the treadmill. The speed of the treadmill was adjusted by the motor and was calibrated. Visual and auditory stimuli were generated using MATLAB. The visual stimulation was delivered to the mice by a video screen placed ∼30 cm from the contralateral eye relative to the virus-expressing hemisphere. The visual and auditory stimuli were synchronized with the imaging using an Arduino board. Where indicated, drugs were injected i.p. 30 minutes before the experiment.

### Immunohistochemistry and immunofluorescence

Immunohistochemistry was performed as previously described^20^. In brief, the mice were anesthetized, the heart was perfused, and the brain was extracted and post-fixed overnight in 4%paraformaldehyde. The fixed brains were then transferred into a PBS-azide solution, and a vibratome was used to cut 45-μm sections. After slicing, free-floating sections were rinsed in washing buffer (PBS containing 0.15% Triton X-100), pre-incubated in 1% hydrogen for 30 minutes, and then rinsed in washing buffer. Sections were blocked for 1 h in washing buffer containing 5% (w/v) bovine serum albumin and 5% (v/v) normal goat serum. After blocking, the sections were incubated overnight in washing buffer containing the rabbit anti-VAChT antibody (Synaptic Systems, #139103; 1:250) plus 2% normal goat serum. After overnight incubation in primary antibody,the sections were rinsed in washing buffer and then incubated for 1 h in an anti-rabbit biotinylated antibody (Vector Laboratories, #ba-9400; 1:200) in washing buffer containing 2% normal goat serum. The sections were then rinsed in washing buffer and incubated with the VECTASTAIN ABC kit (Vector Laboratories) in accordance with the manufacturer’s instructions. The substrate diaminobenzidine (Vector Laboratories) was added as a chromogen, and the sections where counterstained with 0.5% methyl green solution. The sections were then cleared in xylene and mounted on glass slides. For immunofluorescence, slices were prepared as described above and incubated in Tris-buffered saline (TBS) containing 1.2% Triton X-100 for 20 minutes. The sections were rinsed with TBS and blocked for 1 h in TBS containing 5% (v/v) normal goat serum. After blocking, the sections were rinsed twice with TBS,and then incubated for 24 h at 4°C with chicken anti-GFP (Abcam, #ab13970, 1:500) in TBS containing 0.2% Triton X-100 and 2%normal goat serum. After 24 hours, the sections were washed twice for 10 min each in TBS. The sections were then incubated for 1 h in Alexa 488 goat anti-chicken antibody (Life Technologies, #A11039; 1:500) in TBS containing 0.2% Triton X-100 and 2% normal goat serum. The sections were washed twice in TBS for 10 minutes, and then incubated in Hoechst 33342 (ThermoFischer, #H3570;1:500) to counterstain the nuclei. The EVOS FL Auto 2 Cell Imaging System (Invitrogen) was used to visualize the sections.

### Statistical analysis

Except where indicated otherwise, all summary data are reported as the mean ± s.e.m. The signal-to-noise ratio (SNR) was calculated as the peak response divided by the standard deviation of the baseline fluorescence. Group differences were analyzed using the Student’s *t*-test, and differences with a *p*-value <0.05 were considered significant.

## Data and Software Availability

The plasmid pAAV-hsyn-ACh3.0 and pAAV-hsyn-DIO-ACh3.0 have been deposited to addgene (#121922 and #121933) and will be available when published. The custom-written MATLAB, Arduino, and TDT programs will be provided upon request.

## Acknowledgments

We thank Y. Rao for generously sharing a two-photon microscope and X. Lei for providing support of the Opera Phenix High-content Screening System at PKU-CLS. We thank Dr. W. Inoue in the University of Western Ontario for kindly sharing the Multi Conditioning System foot-shocker. Work at the University of Western Ontario (L.S.M., V.F.P, T.B, M.A.M.P.) was supported by the First Research Excellence Fund (CFREF)-BrainsCAN and CIHR (PJT 159781). O.K. was supported by a OGS PhD Fellowship. This work was supported by the General Research Program of National Natural Science Foundation of China (project 31671118), the NIH BRAIN Initiative (grant U01NS103558), the Beijing Brian Initiative of Beijing Municipal Science & Technology Commission (Z181100001518004), the Junior Thousand Talent Program of China, and by grants from the Peking-Tsinghua Center for Life Sciences and the State Key Laboratory of Membrane Biology at Peking University School of Life Science (all to Y.L. L).

## Author contributions

M.J. and Y.L.L. conceived the project. Y.X.L., S.Z., T.Q. and M.J. screened the candidate ACh sensors and characterized the sensors in cultured cells and brain slices. Z.J.Z. and K.T. performed the experiments with transgenic flies. W.P. performed the fiber-photometry recordings of ACh signals during the sleep-wake cycle under the supervision of M.X. P.H. performed the fiber-photometry recordings of foot-shock–induced ACh signals in the BLA under the supervision of H.L. M.S. and O.K. performed the recordings in VAChT^-/-^ mice under the supervision of L.M.S, V.F.P, T.M and M.A.M.P. S.P. performed the luciferase complementation assay. M.J. and R.W. performed the ACh imaging experiments in the visual cortex using miniature two-photon microscopy under the supervision of L.C. and H.C. All authors contributed to the data analysis. M.J. and Y.L.L. wrote the manuscript with input from all other authors.

---

**Figure S1:**
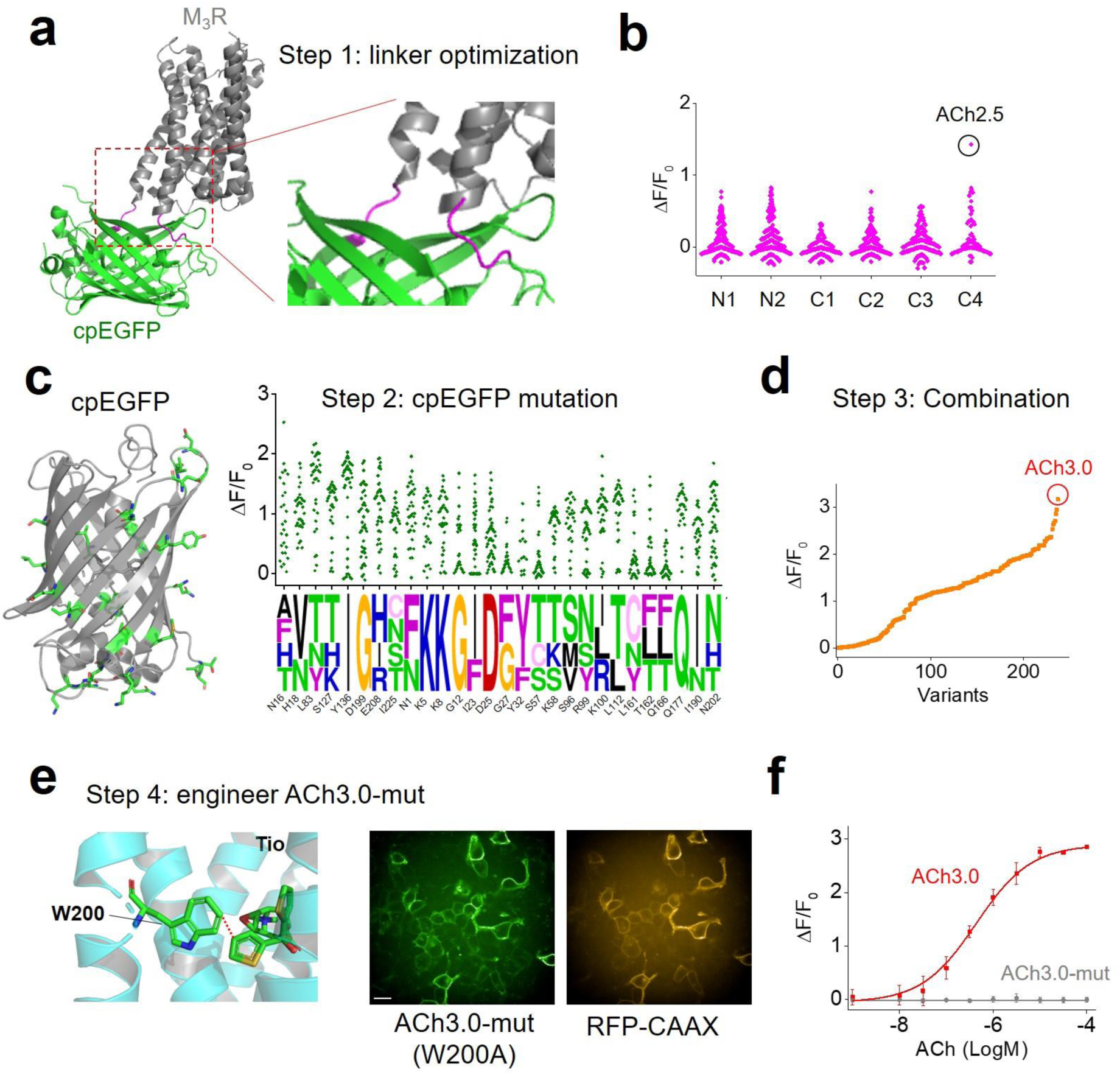
The engineering process leading to the GRAB_ACh3.0_ sensor (related to Fig. 1). **A:** Schematic illustration depicting the predicted structure of the generic GRAB_ACh_ sensor, with the linker region between the receptor (M_3_R) and cpEGFP magnified at the right and shown in magenta. **B:** Site-directed mutagenesis of residues in the N and C termini of the linker region. The numbers indicate amino acid positions relative to the start/end of the cpEGFP sequence. The candidate with the best response is shown in a black circle and was called ACh2.5; this candidate was used for further engineering steps. **C:** Left: crystal structure of the cpEGFP moiety in the ACh3.0 sensor; targeted residues for mutagenesis screening are indicated in green. Right, the fluorescence response of the indicated mutant candidate sensors is shown on top, with the sequences of the best-performing candidates on the bottom; the relative size of each letter reflects the probability of that amino acid in the sequence. **D:** The fluorescence response of each candidate ACh sensor with combined mutations from the best-performing sites in the linker and cpEGFP. Each point is calculated from the average of >100 cells. **E and F:** The W200A mutation in the ligand-binding pocket of M_3_R results in the ACh3.0-mut sensor, which is expressed robustly in HEK293T cells and traffics to the membrane (shown by co-localization with membrane-targeted RFP-CAAX), but produces no detectable fluorescence change in response to ACh; also shown is the ACh-induced response in cells expressing ACh3.0 (n=3 wells for each point, with each well averaging >100 cells). Scale bar represents 10 μm.

**Figure S2:**
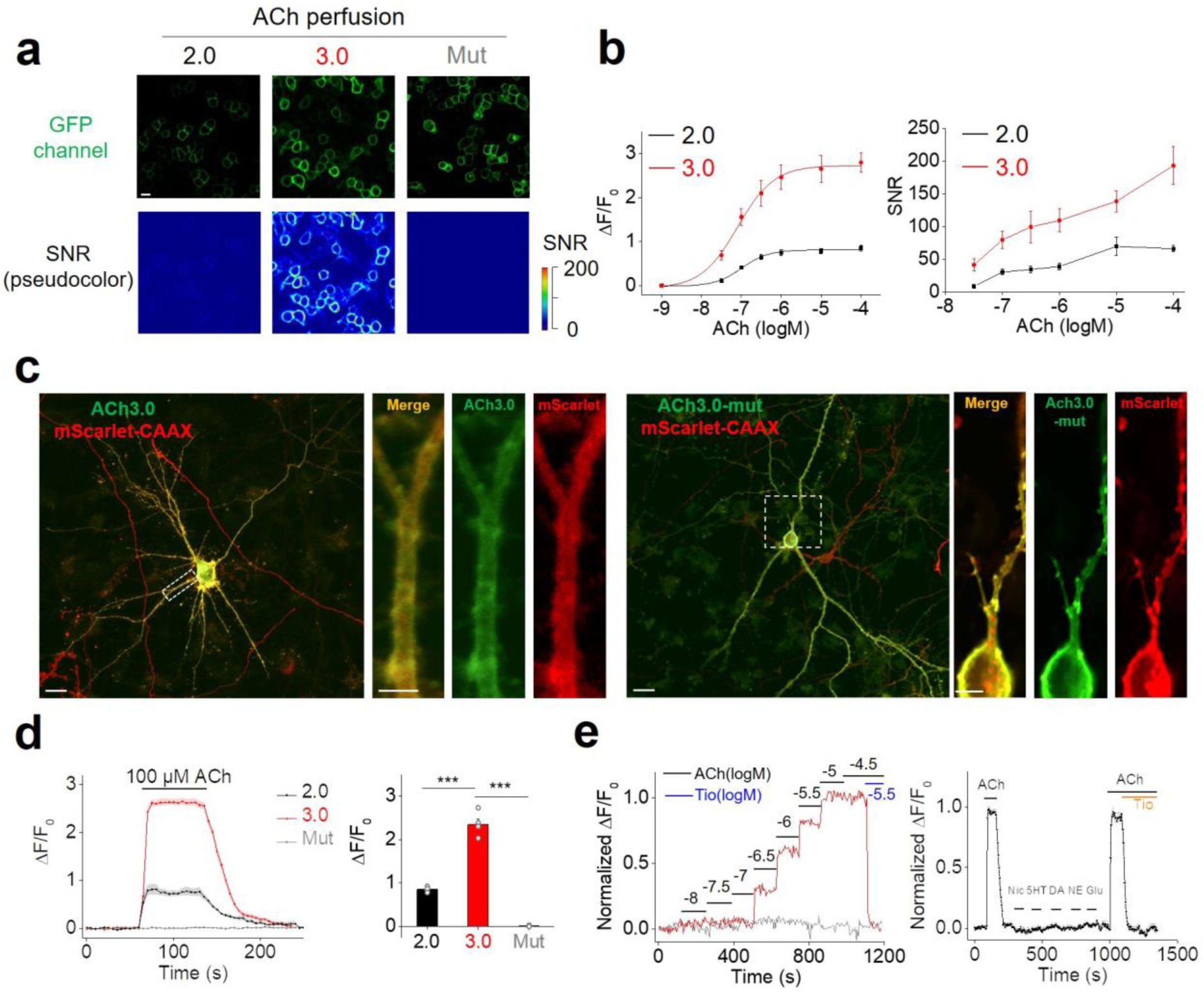
Characterization of ACh2.0 and ACh3.0 (related to Figure 1). **A:** The fluorescence response of ACh2.0, ACh3.0, and ACh3.0-mut to 100 μM ACh. The fluorescence images are shown on top, and corresponding pseudocolor images representing the signal-to-noise ratio (SNR) are shown at the bottom. Scale bars represent 10 μm. **B:** The peak fluorescence response (ΔF/F_0_, left) and SNR (right) of ACh2.0 (black) and ACh3.0 (red) were measured with the indicated concentrations of ACh; n=8 and 7 cells for ACh2.0 and ACh3.0, respectively. **C:** Example fluorescence images of ACh3.0 (left) and ACh3.0-mut (right) expressed in cultured neurons. Membrane-targeted mScarlet-CAAX was co-expressed and used to confirm expression at the plasma membrane. Scale bars represent 10 μm in the original image and 5 μm in the magnified images. **D:** Representative traces (left) and group summary (right) of the fluorescence response of ACh2.0, ACh3.0, and ACh3.0-mut expressed in cultured neurons; where indicated, 100 μM ACh was applied to the cells (n=4, 5, and 7 neurons for ACh2.0, ACh3.0, and ACh3.0-mut, respectively). **E:** Left, representative traces of the normalized fluorescence change in ACh3.0 (red) and ACh3.0-mut (gray) in response to application of the indicated concentrations of ACh. Note that the ACh-induced fluorescence response in ACh3.0 was blocked by the M_3_R antagonist tiotropium (Tio, 3 μM). Right, representative trace of the normalized fluorescence change in ACh3.0 in response to indicated compounds. ACh: 100 μM; nicotine (Nic): 50 μM; 5-HT: 1 μM; norepinephrine (NE): 10 μM; dopamine (DA): 20 μM; glutamate (Glu): 10 μM; and Tio: 2 μM. \*\*\**p*<0.001.

**Fig. S3:**
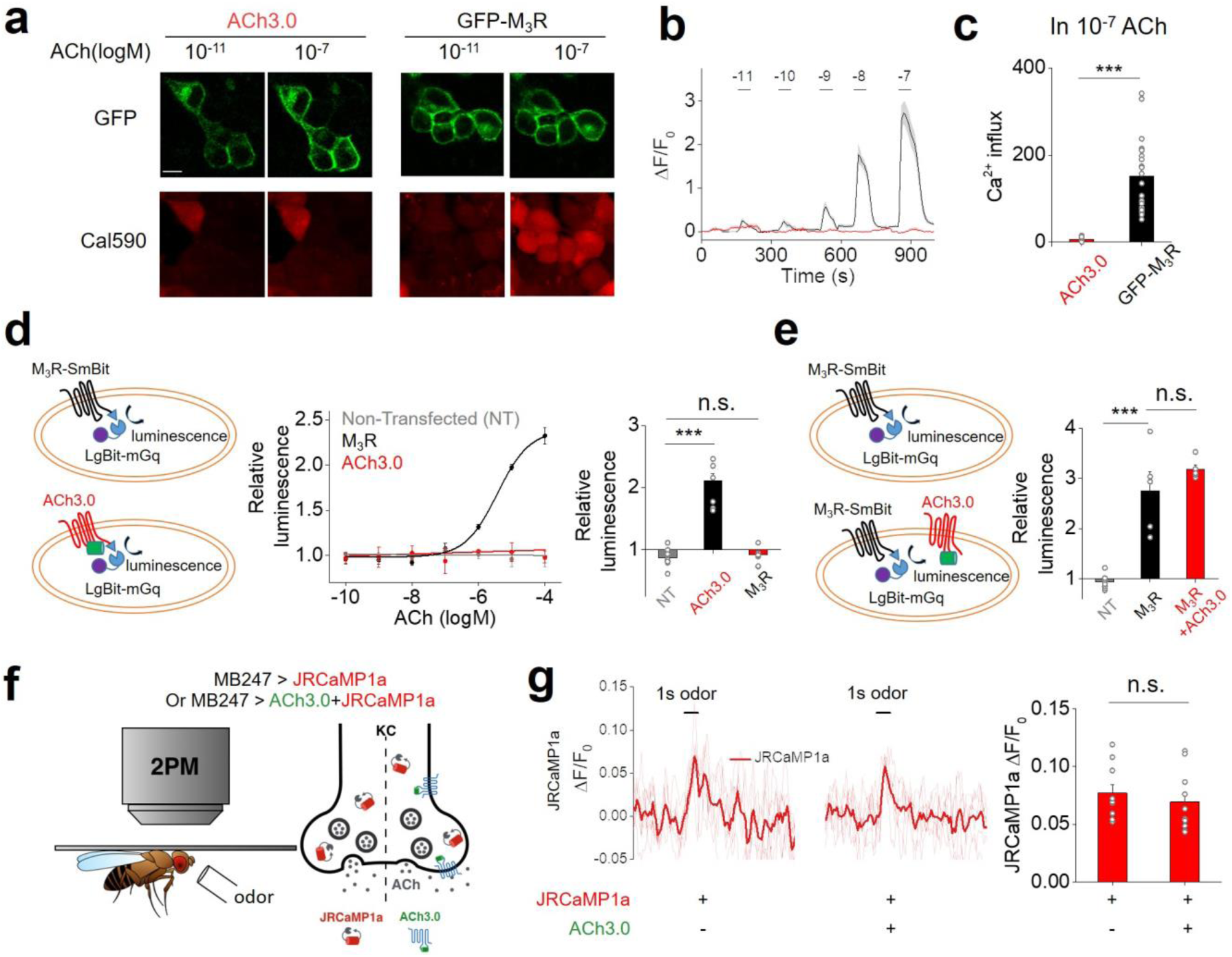
The GRAB_ACh3.0_ sensor produces negligible downstream signaling (related to Figures 1 and 2). **A-C:** Cells expressing either a GFP-tagged M_3_R construct or ACh3.0 were loaded with the red calcium dye Cal590 (A), and the change in Cal590 fluorescence was measured in response to various concentrations of ACh (B). The calcium influx is calculated as the integration of Cal590 fluorescent signal (ΔF/F_0_) to ACh application. The group summary data for calcium influx measured in response to 0.1 μM ACh are shown in panel C; n=21 and 15 cells for GFP-M_3_R and ACh3.0, respectively. **D:** Left, cartoon illustrating the experimental design of the luciferase complementation assay, in which cells expressed M_3_R-SmBit or ACh3.0-SmBit together with LgBit-mGq. Middle, the luminescence signal measured in non-transfected cells (NT), cells expressing ACh3.0-SmBit, and cells expressing M_3_R-SmBit in response to application of the indicated concentrations of ACh, normalized to the signal measured in control buffer-treated cells. Right, group summary of the luminescence signal measured in response to 100 μM ACh; n=9 wells per group, with each group averaging >100 cells. **E:** Similar to D, except the luminescence signal was measured in cells expressing M_3_R-SmBit and cells expressing both M_3_R-SmBit and ACh3.0. The group summary at the right shows the luminescence signal in response to 100 μM ACh; n=5-8 wells per group, with each group averaging >100 cells. **F:** Schematic cartoon depicting two-photon imaging of transgenic flies in response to odorant stimulation. Calcium influx was measured by expressing jRCaMP1a either alone or together with ACh3.0 in the Kenyon cells (KC) in the mushroom body. **G:** Representative fluorescence traces (left) and group summary (right) of jRCaMP1a fluorescence measured in response to odorant application in flies expressing jRCaMP1a either alone or together with ACh3.0; n=10 flies per group. Scale bar represents 10 μm. \*\*\**p*<0.001 and n.s., not significant.

**Figure S4:**
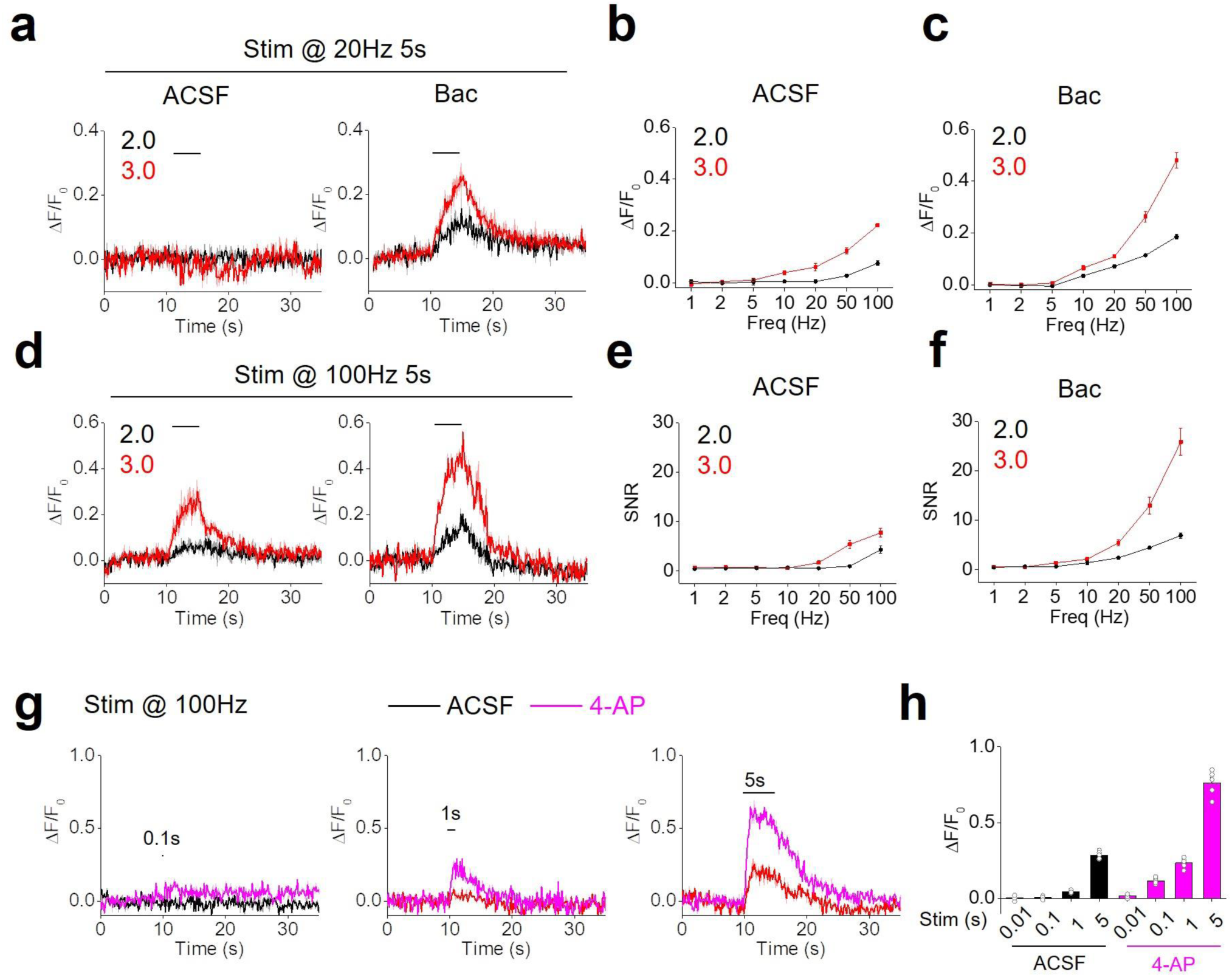
Probing endogenous ACh release in mouse brain slices (related to Figure 2). **A-F:** Representative fluorescence traces (A and D) and group summary of the fluorescence change (ΔF/F_0_ and SNR) in neurons expressing either ACh2.0 or ACh3.0 in response to electrical stimulation in MHb-IPN brain slices. The slices were bathed in either ACSF or 2 μM baclofen (Bac). N=5 slices from 3 mice for ACh2.0, and n=10 slices from 7 mice for ACh3.0. **G-H:** The representative fluorescence traces and group data of ACh3.0-expressing neurons to 100-Hz electrical stimulation with different stimulation times in MHb-IPN brain slices. The response in either ACSF or 100 μM 4-AP was measured and summarized; n=5 slices from 5 mice.

**Figure S5:**
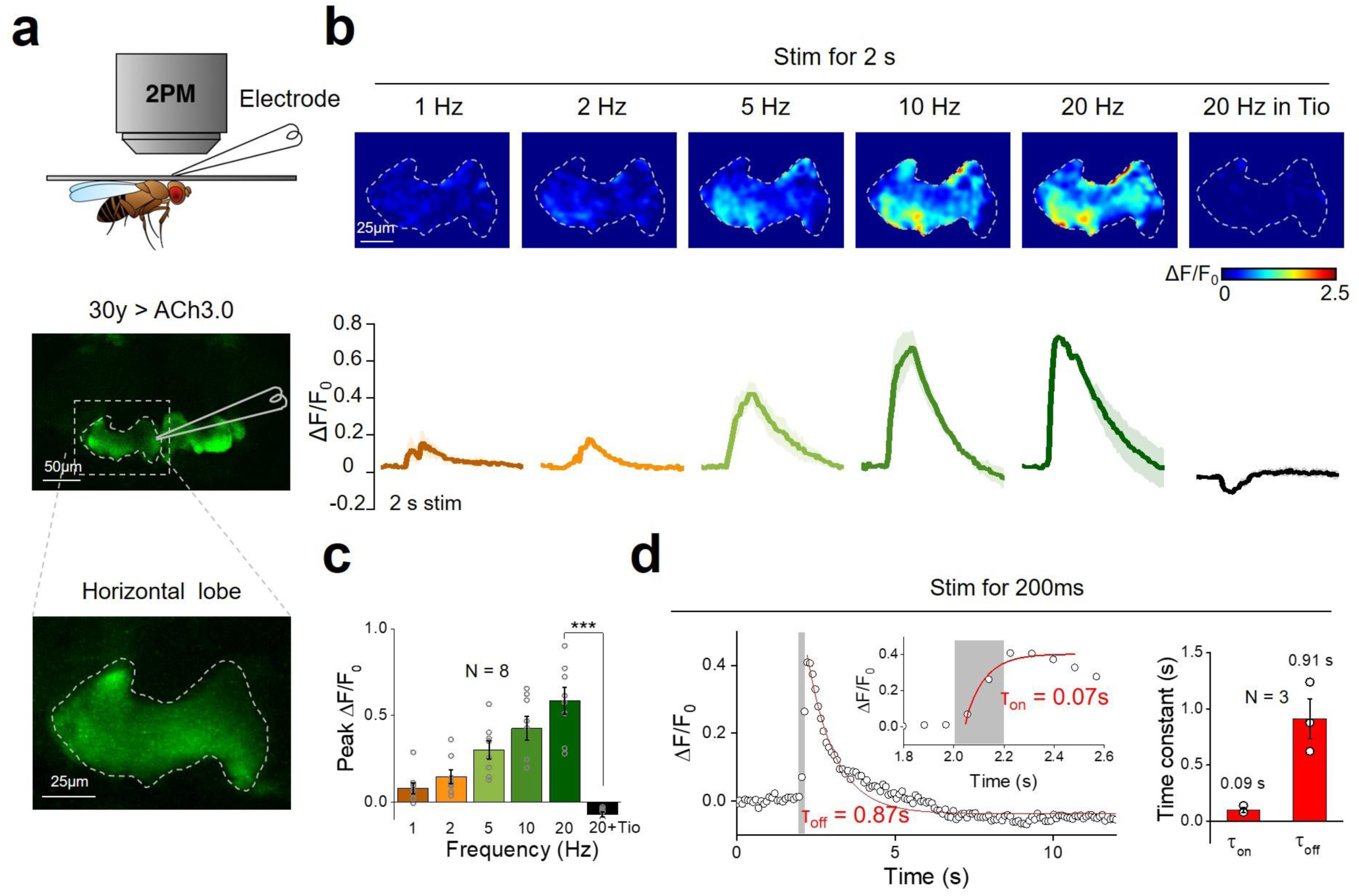
Monitoring *in vivo* ACh release induced by electrical stimulation in *Drosophila* (related to Figure 2). **A:** Schematic illustration depicting the experiment in which a transgenic fly expressing ACh3.0 in the KC cells in the mushroom body were placed under a two-photon microscope, and a glass electrode was placed near the mushroom body and used to deliver electrical stimuli. **B:** Pseudocolor images (top) and representative traces (bottom) of the fluorescence change in ACh3.0 in response to 2 s of electrical stimulation at the indicated frequencies. Where indicated, the M_3_R antagonist tiotropium (Tio, 10 μM) was applied to the bath solution. **C:** Group summary of the data shown in panel B; n=8 flies. **D:** ACh3.0 fluorescence was measured before and after a brief (200 ms) electrical stimulation, and the rising and decay phases were fitted with a single-exponential function; the time constants are indicated and are summarized on the right; n=3 flies. \*\*\**p*<0.001.

**Figure S6:**
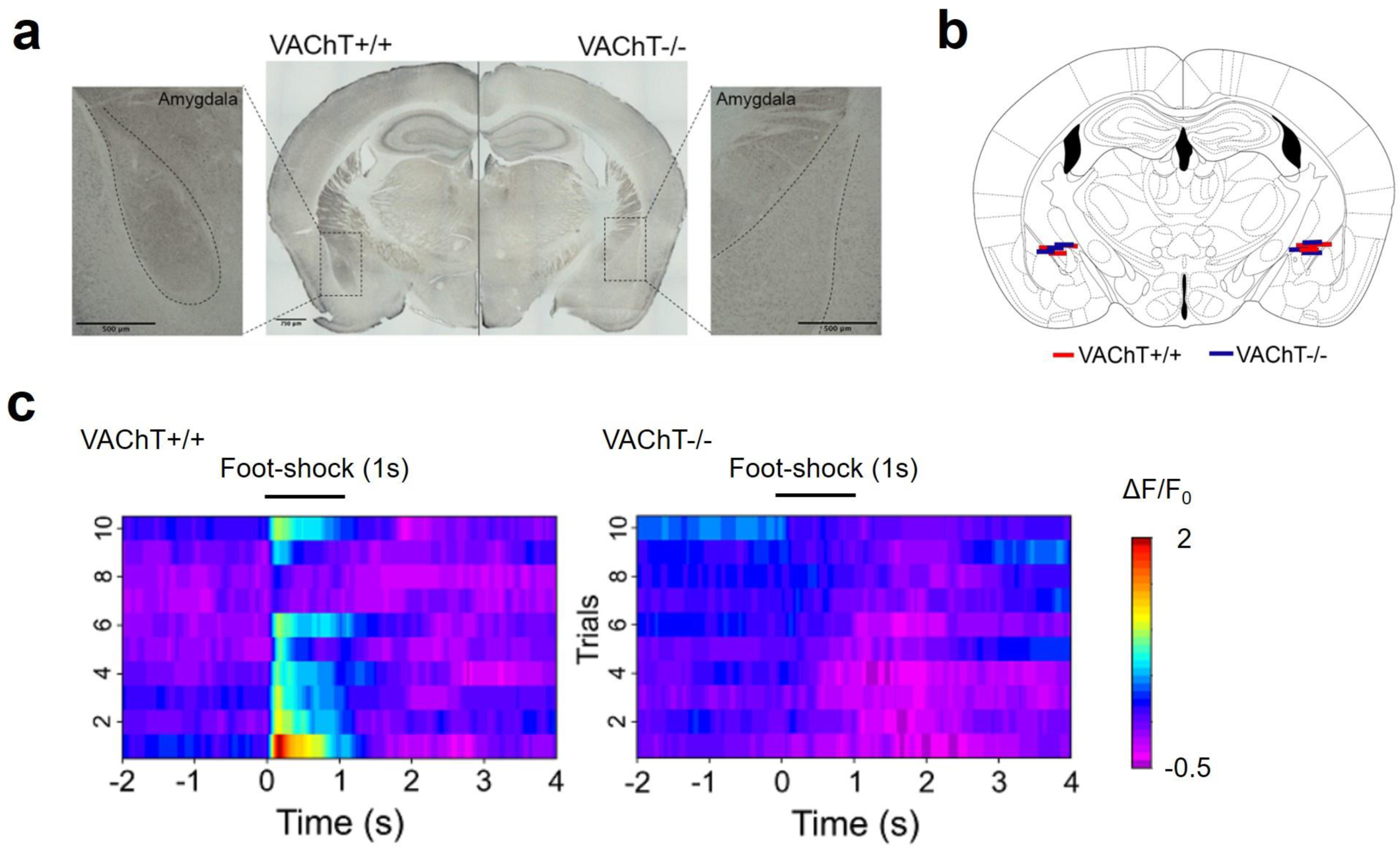
Monitoring endogenous cholinergic signals in mice *in vivo* (related to Figure 3). **A:** VAChT immunohistochemistry was performed in coronal mouse sections obtained from a control (VAChT^+/+^) mouse (left) and a VAChT^-/-^ mouse (right). The insets show magnified views of cholinergic terminals in the basolateral amygdala (BLA). Scale bars represent 500 μm. **B:** Diagram summarizing the location of the optic fiber terminals in the BLA of VAChT^+/+^ (red) and VAChT^-/-^ mice (blue); n=6 mice per group. **C:** Pseudocolor image showing the change in ACh3.0 fluorescence measured in the BLA of VAChT^+/+^ mice (left) and VAChT^-/-^ mice (right) in response to a 1-s foot-shock; ten consecutive trials are shown.

